# Defined media reveals the essential role of lipid scavenging to support cancer cell proliferation

**DOI:** 10.1101/2025.02.12.637975

**Authors:** Oliver J. Newsom, Lucas B. Sullivan

## Abstract

Fetal bovine serum (FBS) is a nearly ubiquitous, yet undefined additive in mammalian cell culture media whose functional contributions to promoting cell proliferation remain poorly understood. Efforts to replace serum supplementation in culture media have been hindered by an incomplete understanding of the environmental requirements fulfilled by FBS in culture. Here, we use a combination of live-cell imaging and liquid chromatography-mass spectrometry to elucidate the role of serum in supporting proliferation. We show that serum provides consumed factors that enable proliferation and demonstrate that the serum metal and lipid components are crucial to sustaining proliferation in culture. Importantly, despite access to a wide range of lipid classes, albumin-bound lipids are the primary species consumed during cancer cell proliferation. Furthermore, we find that combinations of the additive ITS, containing necessary metals, and albumin-associated lipid classes are sufficient to replace FBS in culture media. We show that serum-free media enables sensitive quantification of lipid consumption dynamics during cell proliferation, which indicate that fatty acids (FA) are consumed through a mass-action mechanism, with minimal competition from other lipid classes. Finally, we find that pharmacologic disruption of FA activation and incorporation into the cellular lipidome reduces uptake from the environment and impairs cell proliferation. This work therefore identifies metabolic contributions of serum in cell culture settings and provides a framework for building cell culture systems that sustain cell proliferation without the variable and undefined contributions of FBS.

## Introduction

Since the origins of cell culture, animal serum has been an essential additive for supporting robust and sustained growth of most cell lines (T. Yao and Asayama 2017; Burrows 1910). Among the various serum additives used in culture media, fetal bovine serum (FBS) is the most widely used in biomedical research, primarily attributed for its roles in providing a complex mix of growth factors, lipids, and other nutrients (Yao and Asayama 2017; Puck, Cieciura, and Robinson 1958). However, despite the ubiquitous use of FBS, its fundamentally undefined nature poses a challenge for scientific understanding, since the many biologically active components cannot be readily uncoupled for study using traditional culture media formulations. In addition, the inherent batch-to-batch variability of FBS can influence experimental outcomes and thus contribute to reproducibility issues (Koch et al. 2021; Liu et al. 2023). Aside from its biological influence, FBS also has the potential to introduce zoonotic pathogens into cell culture systems, resulting in safety concerns, particularly in the context of biomanufacturing (Chou et al. 2015). Finally, it is unclear whether fetal bovine fluid is relevant for studying post-natal human disease, as its composition may inappropriately influence biological phenotypes and inadequately model the in vivo environment. These issues therefore highlight the need to develop serum-free alternatives for cell culture.

A major hurdle to excluding FBS from cell culture media is that serum components required to sustain cell proliferation remain poorly defined. For instance, although growth factors are a well-known constituent of FBS, the extent to which specific serum-derived hormones drive cell proliferation remains uncertain (Barnes and Sato 1980). In the case of cancer cell culture, the requirement for fetal serum rich in growth factors is questionable, as a hallmark of cancer is constitutive activation of growth factor signaling pathways in the absence of exogenous cues (Hanahan and Weinberg 2011). Moreover, fetal growth factors may impair our understanding of oncogenic signaling in cancer by masking phenotypic effects of alterations to cancer-associated growth factor pathways. Alternatively, FBS may support cancer cell proliferation by providing small molecule nutrients, including lipids, soluble metabolites, vitamins, and trace metals. Indeed, several studies have found that modifying the relative proportion of these molecules in media through various filtering, extraction, and formulation strategies can influence genetic and metabolic dependencies of proliferating cancer cells (Abbott et al. 2023; Rossiter et al. 2021; Svensson et al. 2016; Wu et al. 2024). The potential nutritional contributions of FBS are also implied when considering that some biologically important vitamins (e.g. biotin and cobalamin) and metals (e.g. iron, zinc, and copper) are absent in many commonly used cell culture media formulations (Rossiter et al. 2021; McKee and Komarova 2017). Indeed, culture media additives that support proliferation in serum restricted conditions often include trace metals or the metal carrier protein transferrin (Yao and Asayama 2017; Ham 1965). Nonetheless, the development of a serum-free media that enables robust proliferation across a broad range of cell lines remains a challenge since the role of serum remains unclear. Collectively, these issues underscore the need for a minimal media capable of supporting cell proliferation without the addition of serum, which would facilitate investigations into the environmental requirements for proliferation, including the metabolic and signaling factors that are otherwise inextricably fulfilled by the numerous serum components.

To explore environmental dependencies of serum for cell proliferation in culture, we combined live cell imaging with liquid chromatography-mass spectrometry (LC-MS) to investigate the role of FBS components— particularly serum lipids—in supporting cancer cell proliferation. Despite the broad range of lipid species in FBS, our findings reveal that cells selectively consume albumin-bound lipid classes during proliferation. Furthermore, combining these lipid classes with cell culture additives that provide micronutrients is sufficient to function as an effective FBS replacement, enabling the uncoupling of metabolic variables that are normally provided by FBS. Our results underscore the critical role of lipids in supporting cell growth and provide a platform to investigate lipid metabolism and other contributions of serum in a defined environment.

## Materials and Methods

### Reagents

Unlabeled oleic acid (O1008) and palmitic acid (P0500) used in proliferation experiments were purchased from Sigma. U-13C oleic acid (CLM-460), U-13C palmitate (CLM-409-0.1), and d11-arachidonic acid (10006758) isotope standards were obtained from Cambridge Isotopes. 5-13C oleic acid (9004089), 4-13C palmitate (30550), LPC 18:1 (20959), and LPC 16:0 (10172) used in consumption assays were purchased from Cayman Chemical. 18:1 LPE (846725P), LPE 16:0 (856705P), 18:1-d7 LPC (791643), 18:1-d7 LPE (791644), and lipids used to generate calibration curves SPLASH® LIPIDOMIX® Mass Spec Standard (330707), LightSPLASH™ LIPIDOMIX® Quantitative MS Primary Standard (330732), were purchased from Avanti Polar Lipids. The chemical inhibitors used in this study GSK2194069 (SML1259) and triacsin C (2472) was purchased from Sigma and Tocris, respectively. FBS lots used for serum lipid comparison were purchased from Cytiva (SH30396.03; Lot No: AH30469640, and AK30775909), Gibco (A52567-01; Lot No: M3009140RP), and Corning (35-077-CV; Lot No:22023001).

### Cell Culture

Cell lines were acquired from ATCC (143B, H1299, HCT116, HT1080, A549, Jurkat), JCRB Cell Bank (OCUG1), or as a gift from Dr. Supriya Saha, Fred Hutch (CCLP1, SSP25). H1299 NucRFP cells were previously generated, as described in (Davidsen et al. 2024). Cell identities were confirmed by satellite tandem repeat profiling and cells were tested and found to be free from mycoplasma (MycoProbe, R&D Systems). Cells were maintained in Dulbecco’s Modified Eagle’s Medium (DMEM) (Gibco, 50–003-PB) supplemented with 3.7 g/L sodium bicarbonate (Sigma-Aldrich, S6297), 10% FBS (Cytiva, SH30396.03) and 1% penicillin-streptomycin solution (Sigma, P4333). Cells were incubated in a humidified incubator at 37 °C with 5% CO_2_.

### Reagent preparations

Numerous cell culture additives containing Insulin-Transferrin-Selenium (ITS) are currently available, often containing supplements beyond its namesake additives, resulting in heterogeneous ITS formulation offerings that are sometimes referred to as “ITS-plus.” The ITS mixture used here was that of Insulin-Transferrin-Selenium Solution Plus (100X), Animal Free (GenDEPOT, CA202), which was used as purchased or made in-house by assembling its components (or subsets thereof, as described) into a 100x stock solution containing 0.5 g/L recombinant insulin (Sigma I9278), 0.6 g/L recombinant transferrin (Sigma, T8158), 0.67 mg/L of sodium selenite (Sigma, 214485), 10 mg/L ethanolamine (Sigma, E0135), 2.04 mg/L ferric citrate (Sigma, F3388), 863 μg/L of zinc sulfate (Sigma, 83265) and 1.6 μg/L of copper(II) sulfate (Sigma, C8027) in water. These solutions were used as a spike-in supplement at 1% into culture media. Lipid mixes were made by dissolving lyophilized lipids in 100% MeOH to a final concentration of 100 mM and lipid solutions were aliquoted and stored in glass vials at −20 °C in glass. Lipid solutions were then diluted to a final concentration of 200 μM in DMEM media containing 50 μM Bovine Serum Albumin Fraction V, heat shock, fatty acid free (Sigma, 3117057001) and placed in a flask and shaken at 37 ° C for 1 hour. 200 μM lipid solutions were then diluted in DMEM to the final working concentration. Unless otherwise noted, lipid mixes were combined at a 95:5 molar-ratio of oleate:palmitate FA chains for both FA mix and LPC mix, while LPE lipid mix was a 60:40 molar-ratio of oleate:palmitate FA chains.

### Incucyte/proliferation experiments

H1299 NucRFP cells were trypsinized (Corning, 25-051-CI), resuspended in media, counted (Beckman Coulter Counter Multisizer 4 or Nexcelom Auto T4 Cellometer), and seeded overnight onto 24-well dishes (Thermo, 142475) with an initial seeding density of 5,000 cells/well. After overnight incubation, wells were washed 3 times in phosphate-buffered-saline (PBS) and 1 ml of treatment media was added. Experiments were conducted in DMEM (Corning 50-013-PB) without pyruvate, supplemented with 3.7 g/L sodium bicarbonate (Sigma, S5761) and 1% penicillin-streptomycin solution (Fisher, 15-140-163) and 1 mM sodium pyruvate (Sigma-Aldrich, P8574) and supplemented with the indicated treatments. Plates were imaged in real time using the IncuCyte S3, at 20X magnification with a 400 ms exposure for the red channel and set to capture images every 6 hours. For standard proliferation assays, initial and final cell counts were collected to calculate proliferation rates.

### Consumption experiments

Cells were seeded on 6-well dishes (Corning, 087721B) in DMEM containing 10% FBS. The following day, the wells were washed 3 times in PBS before swapping cells into 2 ml of the indicated media. Plates were placed back in the incubator for 1 hour to allow for the cells and media to equilibrate. For consumption experiments in 10% FBS/DMEM, media was collected at the indicated time and cells were counted. For consumption experiments in the serum replacement media, cell counts were collected from a parallel plate at time 0 and 100 μl of media was collected at 2-hour intervals for 6 hours. At the final time point, cell counts were collected from the well to calculate cell hours and proliferation rates during the assay.

### Lipidomic sample preparations

Media samples were extracted using a single-phase extraction protocol, where 30 μl of media was combined with a 270 μl of 2:1 ratio of ethyl acetate (Thermo, 022912.K2):2-propanol (Sigma, 34863) containing isotopically labeled lipid standards in glass conical vials (Microsolv technology, 9512S-0CV-T-RSD), resulting in a final extraction solvent of 6:3:1 ethyl acetate: 2-propanol: water. Samples were vortexed for 10 minutes and centrifuged at max speed to pellet debris. 200 μl of supernatant was transferred to a clean glass vial before drying down using a refrigerated vacuum concentrator (Fisher, 10269602). Once dry, lipids were resuspended in 100ul of 65:30:5 acetonitrile (Sigma, 34851): 2-propanol: water and transferred to glass autosampler (Fisher, 03-452-330) vials for mass spectrometry analysis. Mass spectrometry runs were analyzed using Skyline software for metabolomics analysis (Valk et al. 2018). Semi-targeted lipidomics analysis of serum lipids was normalized to Avanti SPLASH deuterated standard for each class, whereas targeted consumption assays included standards for each individual lipid.

### Isotope Tracing

H1299 cells were seeded in a 6-well dish at an initial density of 2×10^5^ cells per well. The following day, cells were washed twice with PBS and swapped to DMEM without glucose, glutamine, pyruvate, or phenol red (Sigma, D5030) supplemented with 10% dialyzed FBS (Sigma, F0392), 1% penicillin-streptomycin, 25 mM U-^13^C glucose (Cambridge Isotopes Laboratory, CLM-1396), and 4 mM ^12^C glutamine (Sigma, G5792). GSK2194069 treated cells were supplemented with 200 nM of GSK and all plates were placed back in incubator for 24 hours before extraction.

### Complex lipid analysis

A 2 μL injection of sample (held at 10°C in the autosampler) was made onto a hypersil gold column (1.0×150mm with a 1×10mm guard, at 50°C), and was eluted (from 32% to 97% “B” over a 36-minute total run time) using a multi-step gradient. The composition of the mobile phases used consists of “A” = water:acetonitrile at 60:40 v/v with formic acid at 0.1% and 10mM ammonium formate, and “B” = acetonitrile:IPA at 10:90 v/v containing 10mM ammonium formate with formic acid at 0.1%. Eluted analytes were analyzed with a Q-Exactive HF-X at 120k resolution in the MS1 mode in both positive & negative polarities in the same run, with ddMS2 performed on the top 15 precursors at 30k resolution with stepped NCE energies of 25 and 30 respectively. Identification of lipids was made by accurate mass comparison to an in-house database of standards and validated by comparison of the MS2 spectra to predicted fragmentation patterns. Prior to data collection, the performance of the instrument and chromatographic system were evaluated for retention time consistency and signal intensity using an injection of 1,2-distearoyl-sn-glycero-3-phosphocholine (PC 18:0) solubilized in 50:50 (v/v) mobile phase A and B. Additionally, instrument performance and mass calibration are evaluated throughout the sample run by a daily injection of Lipidomix SPLASH mix (Avanti Polar Lipids) containing 14 deuterium-labeled lipid species.

### Fatty acid analysis

Post extraction, fatty acid samples were re-suspended in acetonitrile/IPA/water 65:30:5 v/v/v, and a 2 μL injection was made onto a Kinetex C8 core-shell column (2.1 × 100 mm, 2.6 μm, and attached 2.1 mm i.d. SecurityGuard Ultra C8 guard column). The autosampler is held at 10°C, while the column is kept at 40° C through the run. The mobile phase composition consisted of “A” = water with acetic acid (at 0.1%) mixed with 5% (by volume) of mobile phase “B”, where “B” = acetonitrile/methanol/0.1% acetic acid (80:15:5 v/v/v). A semi-isocratic gradient is used to elute the analytes off the column, beginning at 20% for 0-1 minute, followed by a rapid increase of “B” to 66%, which is then held for 6.5 minutes, followed by a further increase of “B” to 100%. Finally, a short wash phase is followed by a re-equilibration phase of 7 minutes at starting conditions (20%), with total injection-to-injection times of 22 minutes. Eluted analytes were detected with the Q-Exactive HF-X in MS1 mode over a mass range of 210-600 m/z using negative polarity. Analytes were identified by accurate mass comparison to an in-house database of standards. Prior to analysis and throughout the sample series, instrument performance is evaluated using a mixture (diluted 1:4 using 50:50 mobile phase “A” and “B”) containing 10 monounsaturated fatty acids at 10 ng/μL each (Cayman Chemical). Mass calibration, retention times and peak intensities were monitored to ensure consistent performance. To compensate for contamination from background saturated and mono-unsaturated FAs, multiple volumes of media were extracted to generate a dose response curve, allowing for the calculation of serum-attributable FA content.

### Quantification and statistical analysis

All graphs and statistical analyses were performed in GraphPad Prism 9.0. Technical replicates, defined as parallel biological samples independently treated, collected, and analyzed during the same experiment, are shown. Experiments were verified with independent repetitions showing qualitatively similar results. Details pertaining to all statistical tests can be found in the figure legends.

## Results

### Serum provides essential consumable factors to support cell proliferation

FBS is a key additive to culture media for supporting cell proliferation, yet the mechanism by which serum-restriction impairs cell proliferation remains incompletely understood. Cell proliferation measurements typically utilize initial and end-point cell counts to calculate proliferation rates while assuming a constant doubling time throughout the experiment, which is unlikely to be a reasonable assumption in all settings. For instance, we propose two hypothetical models of proliferation defects that can result in the same final cell count: (1) A rate-limitation phenotype, where restriction of growth factors results in a slowed but constant proliferation rate, or (2) a depletion phenotype, where proliferation rate remains stable initially but decreases once essential serum factors are exhausted (Fig S1A). Given that these two models reflect different biological mechanisms, we hypothesized that understanding the kinetics of proliferation defects during FBS restriction may inform how FBS supports cell proliferation.

To investigate the contribution of FBS to proliferation, we expressed nuclear-localized red fluorescent protein in H1299 non-small cell lung cancer cells (H1299 NucRFP) and used live-cell imaging to track nuclei counts in real-time as a readout of cell proliferation in DMEM media with various FBS concentrations. As expected, FBS was required for sustained cell proliferation, and final cell counts were proportional to media FBS concentration (Fig 1A, left). Interestingly, when comparing the kinetics of cell population growth over time, initial increases in cell counts were similar across all FBS concentrations, however the rate of cell population growth slowed over time proportional to FBS concentrations (Fig 1A, right). When we converted changes in cell count to time-resolved proliferation rates, the kinetics closely matched those predicted for a depletion phenotype (Fig 1B, Fig S1A).

**Figure 1.**
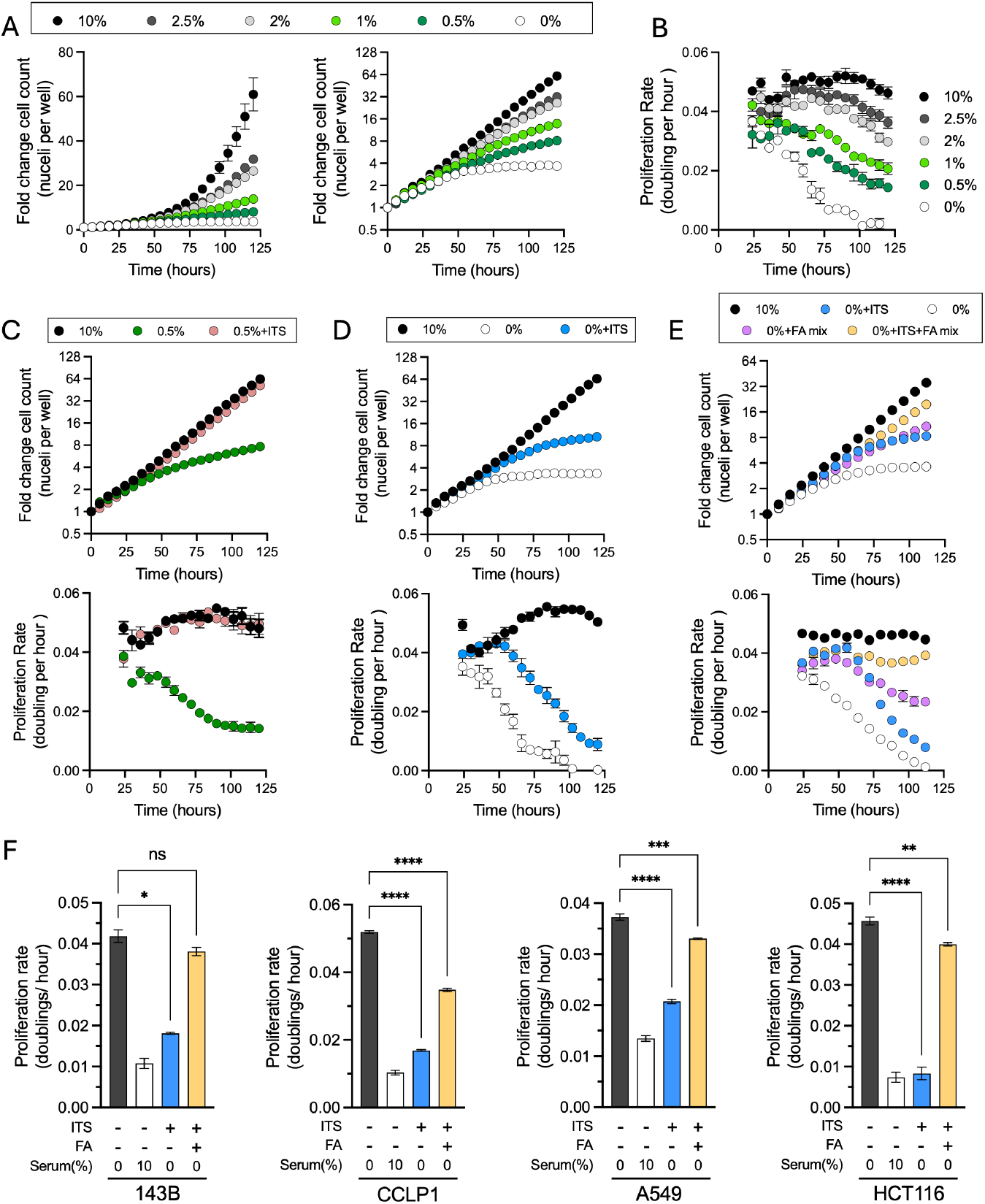
Effects of serum limitation on cancer cell proliferation rates. **(A)** Fold change in H1299 NucRFP cell counts over time when cultured with varying concentrations of FBS. Data are shown on a linear scale (left) and a log2 scale (right). **(B)** Moving average of the proliferation rate (calculated from four adjacent time points) for cell nuclei counts measured in (A). **(C)** H1299 NucRFP cells grown in 10% FBS or 0.5% FBS treated with vehicle or ITS. Fold change in cell counts (top) and moving average of proliferation rate (bottom) **(D)** H1299 NucRFP cells cultured in 10% FBS or in DMEM alone, treated with or without ITS. Fold change in cell counts (top) and moving average of proliferation rate (bottom) **(E)** H1299 NucRFP cells cultured in 10% FBS or in DMEM alone and treated with either a FA mix (100 μM), ITS, or both. Fold change in cell counts (top) and moving average of proliferation rate (bottom) **(F)** Proliferation rates of A549, CCLP1, HCT116, and 143B cells grown in 10% FBS compared to DMEM supplemented with either ITS or a combination of ITS and FA mix, measured by initial/final cell counts. Error bars indicate mean ± SEM (n=3). Abbreviations: ITS, insulin–transferrin– selenium mix that also contains ethanolamine and trace metals; FA, fatty acid mix. Statistical significance was assessed using one-way ANOVA Brown-Forsythe and Welch ANOVA tests (F). ns = not significant, **p < 0.05, **p < 0.01, ***p < 0.001, ****p < 0.0001. 13

Notably, upon slowing, the rate of proliferation decay was comparable in each case of FBS limitation, with the primary difference being that the onset of proliferation defects occurred later in conditions with higher FBS concentrations (Fig 1B). These findings therefore support the hypothesis that FBS restriction impairs cell proliferation by limiting consumable components necessary for sustained proliferation.

Since basal media lacks numerous components of FBS that could support proliferation, we next tested whether supplementation with a defined additive containing insulin, transferrin, selenium, ethanolamine, and biologically relevant metals (ITS, see methods), a mixture shown to shown to support cell proliferation in serum restricted conditions, could similarly restore cell proliferation to these cells during FBS limitation (Bottenstein and Sato 1979; Murakami et al. 1982). Indeed, ITS was sufficient to fully restore the proliferation rate kinetics of H1299 NucRFP cells cultured in 0.5% FBS to match that of cells cultured in 10% FBS (Fig 1C). We next investigated which component(s) of the ITS mixture mediated the proliferation restoration in low FBS conditions and observed that transferrin and metals are the dominant contributors to H1299 NucRFP proliferation upon serum-restriction (Fig S1B). Consistent with the limitation of these micronutrients in low serum conditions, titrating down the concentration of ITS in 0.5% FBS restored the depletion phenotype (Fig S1C). These data therefore indicate that a primary limitation in serum restricted conditions is the depletion of trace metals that are not otherwise provided in the basal media. We next tested whether adding ITS could replace FBS altogether. Although ITS improved the overall number of doublings compared to DMEM alone, these cells still exhibited a decaying proliferation rate indicative of a delayed depletion phenotype (Fig 1D). This suggests that while ITS is beneficial under reduced FBS conditions, presumably by solving the first order deficiency for metals, it cannot replace FBS entirely, as it lacks an additional essential serum component that is required to sustain cell proliferation.

We next explored how other factors in FBS may contribute to cell proliferation of ITS treated cells in basal media. Serum-derived lipids serve as a metabolic resource that are utilized during cell proliferation which can be conditionally essential during metabolic stress (Kamphorst et al. 2013; Svensson et al. 2016; Yao et al. 2016; Li et al. 2022). Thus, we tested the impact of lipid supplementation by adding a fatty acid (FA) mix composed of palmitate and oleate, the two most abundant saturated and mono-unsaturated FA species in serum, conjugated to bovine serum albumin (BSA) (Abdelmagid et al. 2015; Else 2020). Like ITS, FA mix improved overall cell proliferation relative to basal media alone, but cell proliferation rate eventually decayed over time (Fig 1E). Importantly, the co-supplementation of ITS with FA mix in basal media prevented cell proliferation rate decay and was sufficient to maintain exponential growth in the absence of FBS, albeit with a modest rate limitation phenotype compared to cells cultured in 10% FBS (Fig 1E, Fig S1D). We also investigated if this combination could support the proliferation of a diverse group of cancer cell lines, including cells deriving from non-small cell lung cancer (A549), osteosarcoma (143B), cholangiocarcinoma (CCLP-1), and colon cancer (HCT116). Measuring proliferation rates by cell count, we observed that the combination of ITS and FA mix was sufficient to support serum-free cell proliferation in each cell line (Fig 1F). While the addition of FBS to culture media still adds convenience for various cell culture functions, including inactivating trypsin while splitting adherent cells and providing attachment factors that encourage cells to adhere to culture dishes, we found that this FBS replacement formulation can maintain robust cell proliferation in Jurkat suspension cells for greater than four weeks (Fig S2). These findings therefore highlight that the requirement for serum can be largely obviated for cancer cell culture experiments by providing a source of metals and fatty acids, which are otherwise absent from basal media (Fig S1D).

### FBS lipidome and its consumption by proliferating cells

While our data demonstrate that exogenous FAs can fulfill a major role of FBS of providing lipids to support growth in culture, FBS is a heterogeneous mixture that contains a wide repertoire of lipid species and so it remains unclear which serum lipids are normally consumed by cultured cells. Because multiple lipid species may redundantly support proliferation, we next examined the lipid composition of FBS and measured which lipids are consumed by proliferating cells. The serum lipidome includes diverse lipid species which vary in the number of FA chains each lipid contains, lipid head group, as well as the length and saturation of the hydrocarbon chains (Quehenberger and Dennis 2011). To characterize the major lipid classes in FBS and their variability across different FBS lots, we used semi-targeted quantitative LC-MS to profile lipid metabolites in four different FBS lots (Fig 2A). Consistent with prior reports, the most abundant lipid classes included neutral lipids, where sterol esters (SE) were much more abundant than triacylglycerols (TAG) (Quehenberger et al. 2010). Additionally, polar lipids like sphingomyelin (SM), and glycerophospholipids, including phosphatidylethanolamine (PE), phosphatidylcholine (PC), and their precursor lysolipids, lyso-PC (LPC) and lyso-PE (LPE) were also detected in all FBS samples. Importantly, while the detected lipid classes were consistent across FBS lots, the relative and total abundance of lipid classes varied between different lots of FBS. These findings therefore demonstrate the inherent variability of using serum in cell culture settings and highlight the need to explore the roles of FBS-derived lipids in a more controlled system.

**Figure 2.**
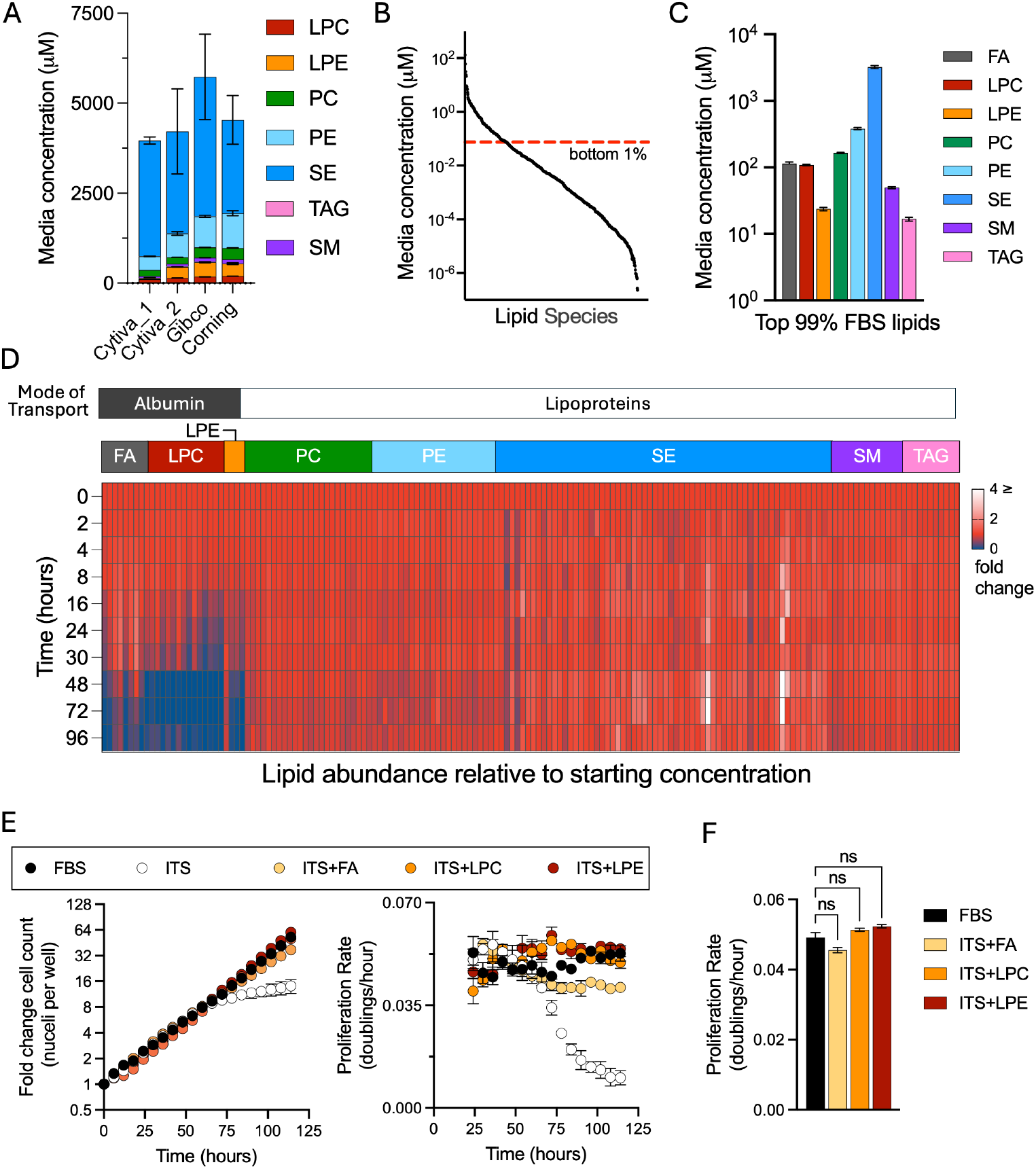
Quantification of lipid depletion during cell proliferation. **(A)** Comparison of lipid profiles across different lots of commercially available FBS. **(B)** All detected lipids ranked from most abundant to least abundant, where lipids which fall below the red line indicate the bottom 1% of total lipids in FBS. **(C)** FBS lipid concentration in 100% FBS used in the remaining experiments. **(D)** Relative concentrations of the top-most abundant lipids normalized to initial concentrations during cell proliferation. **(E)** Proliferation of H1299 cells cultured in either 10% FBS or DMEM supplemented with ITS plus one of the following: LPC, LPE, or FA mix. Fold change in cell counts (left) and moving average of proliferation rate (right). **(F)** Average proliferation rate of cells over the entire assay. Error bars represent mean ± SEM (n = 3). Abbreviations: SE, sterol esters; PE, phosphatidylethanolamine; PC, phosphatidylcholine; SM, sphingolipids; TAG, triacylglycerols; LPC, lysophosphatidylcholine; LPE, lysophosphatidylethanolamine; FA, fatty acids. Statistical significance was assessed using one-way ANOVA Brown-Forsythe and Welch ANOVA tests (F). ns = not significant.

We next sought to determine which serum lipids are consumed by proliferating H1299 NucRFP cells. To focus on the lipids present in FBS at biologically relevant concentrations for proliferation, we ranked the detected lipids from most abundant to least and excluded those that constituted the bottom 1% of the total available FBS lipid pool, narrowing the analysis from 563 species to 163 lipids spread across eight classes, varying in FA chain length and degree of saturation (Fig 2B-C, S3). Among these, SEs were the most abundant lipid class, which were quantified at 3.22 +/-0.098 mM, while TAGs were the least abundant lipid class, at 16.73 +/-0.67 µM. PEs, the most abundant glycerophospholipid detected, and PCs were present at 381.93 +/-8.24 µM and 164.78 +/-2.16 µM, respectively; while SMs were detected at 49.36 +/-1.02 µM. The lipid precursors, LPC and LPE, were detected at lower concentrations of 108.69 +/-1.36 µM and 23.65 +/-0.74 µM. Although background from unlabeled saturated and monounsaturated FAs is a known challenge in lipidomics that affects accurate quantification, we quantified FAs at 113.40+/-7.03 µM, representing only a small fraction of the total serum lipid pool despite sufficiency in supporting proliferation in the serum-substitute media.

To identify which FBS-derived lipids are consumed from the media during proliferation, we collected spent media from cells cultured in 10% FBS containing media at multiple time points and profiled the change in the media lipidome over time. Notably, only a subset of available lipids showed meaningful depletion relative to their starting concentration. Specifically, the lipid precursors containing only a single fatty acid, which include FA, LPC, and LPE were selectively consumed, with > 80% depletion in most species relative to the starting amount over 96 hours (Fig 2D). In contrast, the complex lipids including SE, TAG, PC, PE, and SM exhibited minimal change with <5% depletion in most species during the same period. Notably, the pattern of media lipid consumption was closely associated with the mode of transport in serum. Indeed, highly consumed lipid species including FA, LPC, and LPE, are primarily transported bound to BSA, whereas SE, TAG, PC, PE, and SM are carried in lipoprotein complexes (Law et al. 2019; Dashti et al. 2011; Gomaraschi 2020; van der Vusse 2009; Quehenberger et al. 2010). To determine whether selective albumin-associated lipid consumption is specific to H1299 NucRFP cells or represents a broader phenotype of proliferating cancer cells, we profiled media lipids during growth across multiple cell lines. Notably, depletion patterns were broadly similar across all cell lines tested, with cells exhibiting selective depletion of albumin-bound lipids and minimal consumption of lipoprotein-associated lipids during proliferation (Fig S4). Altogether, the preference for albumin-bound lipids during proliferation across various cells suggests that these lipids fulfill a general metabolic requirement for cancer cell proliferation in culture.

### Consumed lipid species redundantly fulfill the lipid requirements of serum

Our data indicate that FAs can fulfill an essential function of FBS in supporting cell proliferation, yet in the context of FBS-containing media, FAs are consumed alongside LPC and LPE lipid species. We thus sought to determine whether these albumin-associated lipid classes were redundant or distinct in their ability to support cell proliferation. We therefore cultured H1299 NucRFP in serum-free media containing ITS, with or without co-supplementation of individual albumin-associated lipid classes and tracked their effect on proliferation kinetics (Fig 2E). Notably, like the combination of ITS and FA mix, both the LPC and LPE lipid mixes were equally sufficient to fulfill the lipid demand for cell proliferation and prevent the depletion phenotype of cells cultured in the absence of serum. Indeed, the average proliferation rate was not statistically different between the cells grown in 10% FBS and any of these serum-free medias (Fig 2F). These data therefore indicate that albumin-associated lipid classes can redundantly fulfill the lipid requirement for cell proliferation.

### Lipid consumption dynamics are a function of environmental availability

The mechanisms and variables that dictate lipid consumption remain poorly understood. We used our serum-free media formulation as a tool to further explore the factors that influence lipid consumption of proliferating cells, using serum-free media containing FAs and/or LPCs as a lipid source. We first cultured cells across a dilution series of FA or LPC mixes and quantified consumption rates. Similar to 10% FBS, cells grown in serum-free media readily consume exogenous FAs and LPCs (Fig 3 A-B). Notably, the consumption rate for both FA and LPC increased proportionally with media concentration, indicating that lipid consumption is predominantly a function of concentration and cells can scavenge lipids at increased rates when they are more abundant in the environment.

**Figure 3.**
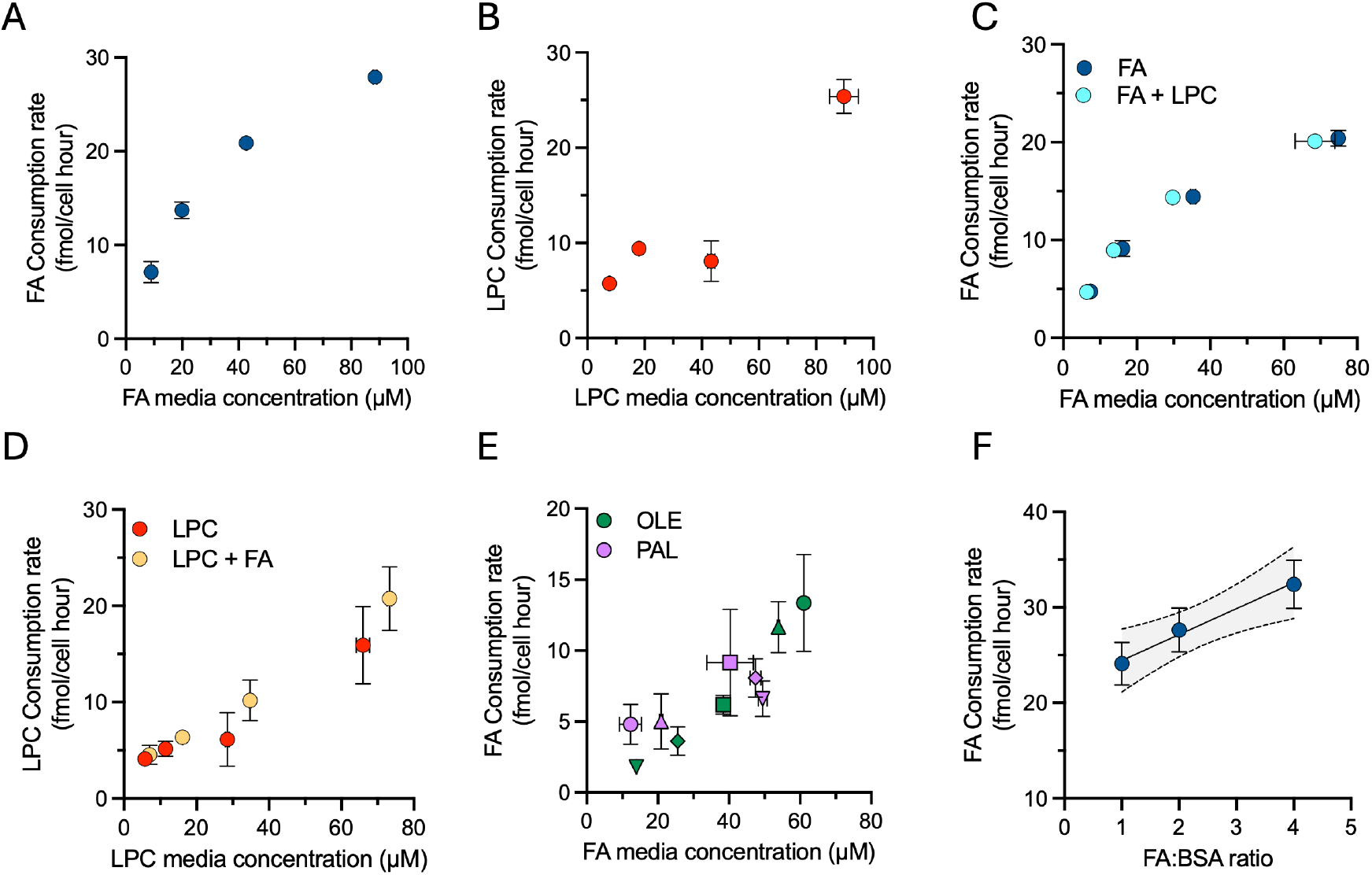
Exploration of lipid scavenging kinetics in serum-free systems. **(A–B)** Consumption rate of total FA (A) and LPC (B) by cells cultured in serum-free media across a concentration series of either lipid mix. **(C–D)** Consumption rates of FA (C) or LPC (D) by cells cultured in serum-free media containing an equimolar mixture of FA and LPC, compared to cells grown in media containing only a serum-free media containing a single lipid class of either FA (C) or LPC (D). **(E)** Consumption rates of palmitate and oleate in cells grown in a fixed concentration of total FA concentration with different molar ratios of oleate-to-palmitate (indicated by matching shapes: ○ 4:1 △ 2:1 □ 1:1 ◊ 1:2 ▽ 1:4). **(F)** Consumption rate of FA in cells cultured at a fixed concentration of FA with varying concentrations of BSA. Error bars represent mean ± SEM (n = 3). Abbreviations: OLE, oleate; PAL, palmitate.

Our data indicate that lipid consumption rates within a lipid class are primarily concentration dependent, but it is unclear if consumption of one class influences consumption of other classes. We next sought to explore how the availability of FA-containing molecules, specifically FAs and LPCs, impacted the consumption of each other. We cultured cells across a dilution series containing an equimolar concentration of FAs and LPCs and compared the consumption rates to those measured when cells were grown in media containing a single lipid class (Fig 3 C-D). Surprisingly, despite sharing BSA as a carrier molecule, FA and LPC uptake was unaffected by the presence of the other lipid class, further supporting non-competitive, concentration-dependent lipid consumption dynamics for both lipid classes. To further explore lipid intrinsic variables affecting lipid consumption, we tested if consumption was selective for FAs based upon chain structure. We thus cultured H1299 nucRFP cells in a fixed concentration of total FAs while varying the molar ratio of palmitate to oleate and quantified consumption of each lipid individually. Interestingly, consumption of each FA followed a similar trend as other uptake variables - showing that, independent of FA chain length or saturation, FA uptake was instead primarily dependent on environmental concentration (Fig 3E). Together, these findings indicate lipid scavenging of FAs and LPCs occurs through a class-independent, mass-action transport mechanism which is unaffected by the FA structure.

Lipids bound to albumin are in dynamic binding equilibrium, and unbound free-FAs are considered the primary form scavenged by cells (Simard, Pillai, and Hamilton 2008). Thus, the availability of lipids for uptake is suggested to be influenced not only by the concentration of lipids in the environment, but also the concentration of albumin, which compete for FA binding with the cell (Spector and Steinberg 1966). We therefore examined how BSA concentration affects FA consumption by culturing cells in media containing a fixed concentration of FA with varying amounts of BSA in the media (Fig 3F). Consistent with previous reports, the consumption of FA from the media scaled proportionally with the FA:BSA ratio, where a higher FA:BSA ratio was associated with increased FA consumption, despite lower BSA available to cells (Simard, Pillai, and Hamilton 2008). These data are consistent with a model in which, as the FA binding sites on BSA approach saturation, the concentration of free-FA increases, leading to proportionally higher lipid consumption. Thus, higher FA:BSA ratios drive increased lipid consumption, supporting the finding that available FA concentration primarily governs consumption rates. Altogether, these data provide insight into the factors governing lipid consumption by proliferating cells and demonstrate the utility of a defined serum substitute media to isolate the variables governing nutrient uptake.

### Lipid scavenging is required for cancer cell proliferation

Metabolic demand is typically a major determinant of nutrient consumption, yet our data indicate that lipid consumption is driven by environmental concentration (Papalazarou and Maddocks 2021). Thus, we next sought to explore whether perturbations to cellular metabolism in FA synthesis or FA activation influenced lipid consumption. We used GSK2194069 (GSK), a fatty acid synthase (FASN) inhibitor, or triacsin C (TriC), a pan inhibitor of the acyl-CoA long-chain (ACSL) family of proteins, which catalyze the condensation of FAs with coenzyme A (Fig 4A) (Tomoda, Igarashi, and Omura 1987; Hardwicke et al. 2014). Using isotope tracing, we confirmed that GSK was effective in preventing *de novo* palmitate synthesis, as evidenced by the loss of isotope incorporation from U-^13^C glucose into palmitate (16:0), the first product of FASN-mediated FA synthesis (Fig 4B). GSK treatment also blocked higher order labeling species of the downstream FA oleate (18:1), but did not abolish the amount of oleate with the M+2 isotopologue (Fig 4B). These data demonstrate on-target inhibition of FASN, as cells maintained the capability to elongate/desaturate unlabeled upstream scavenged FAs to produce oleate. Interestingly, despite substantially impairing FASN activity, GSK had no effect on the cell proliferation of H1299 nucRFP cells cultured in FBS containing media, indicating that *de novo* fatty acid synthesis is not required in these conditions (Fig 4C).

**Figure 4.**
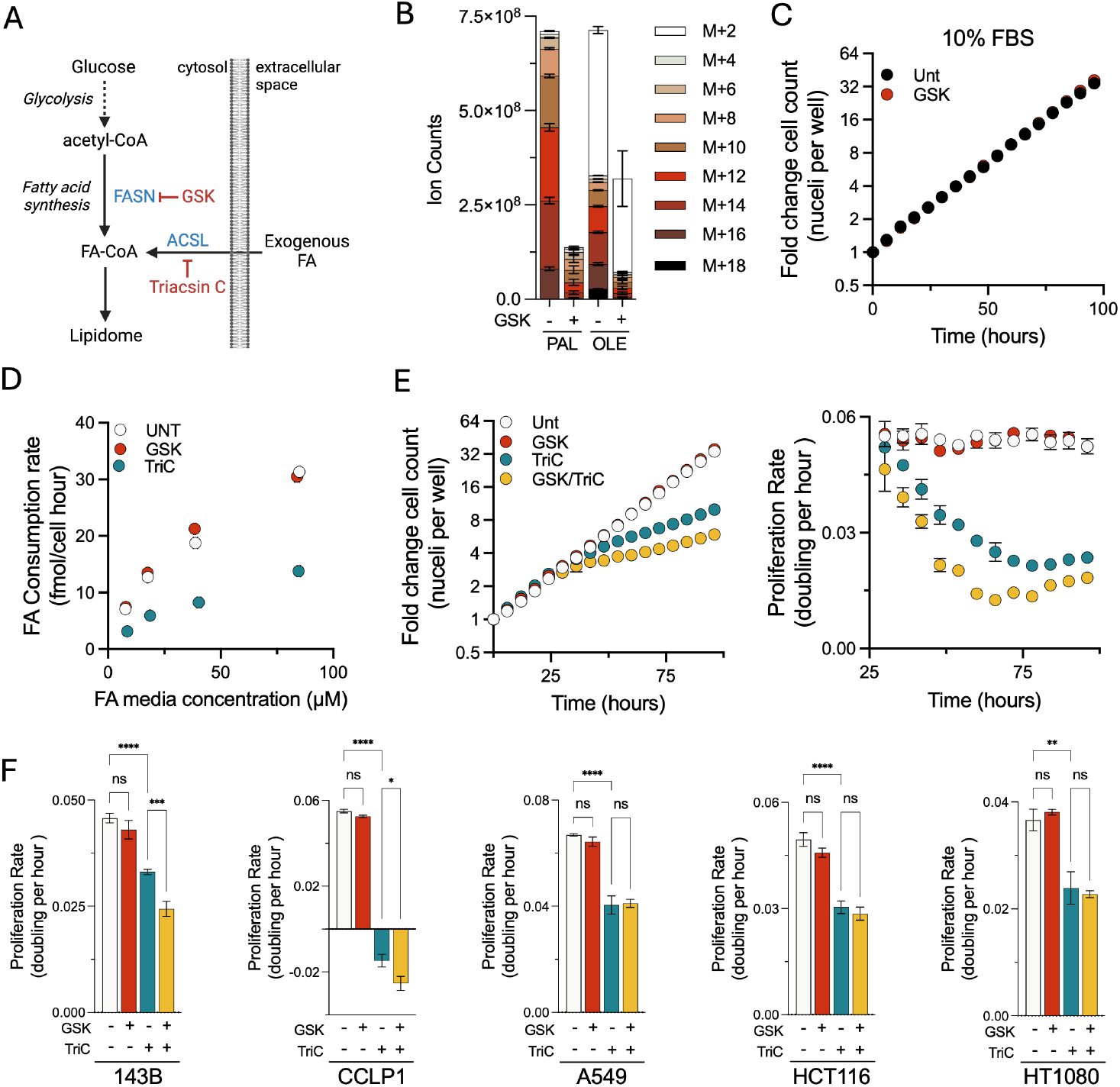
Disruption of fatty acid activation limits lipid scavenging. **(A)** Schematic of FA sources into the cellular lipidome with steps inhibited by GSK and triacsin C. **(B)** Isotopologue distribution of palmitate (C16:0) and oleate (C18:1) in cells cultured in U-13C labeled glucose with or without GSK (200 nM). **(C)** H1299 NucRFP cells grown in 10% FBS containing media treated with or without GSK. **(D)** Consumption rate of FA by cells cultured across a FA dilution series treated with either vehicle, GSK, or triacsin C (4 μM). **(E)** H1299 NucRFP cells grown in serum-free media containing FA treated with either GSK, triacsin C, or both. Fold change in cell counts (top) and moving average of proliferation rate (bottom). **(F)** Average proliferation rate of A549, CCLP1, HCT116, 143B, and HT1080 cells grown in FA containing serum-free media treated with GSK or triacsin. Abbreviations: PAL, palmitate; OLE, oleate. Error bars represent mean ± SEM (n = 3). Statistical significance was assessed using one-way ANOVA Brown-Forsythe and Welch ANOVA tests (F). ns = not significant, *p < 0.05, **p < 0.01, ***p < 0.001, ****p < 0.0001.

FASN impairment had no effect on cell proliferation in standard culture conditions, prompting us to investigate how disruption to lipid metabolism influences FA sourcing. We thus cultured cells across a dilution series of FAs in the presence of GSK or TriC, and quantified FA consumption in the media. Surprisingly, inhibiting FASN-dependent lipid synthesis had no measurable impact on FA consumption, in line with our earlier observation that lipid uptake is primarily determined by lipid availability (Fig 4D). In contrast, TriC treatment resulted in near complete loss of FA consumption from the media at all concentrations, consistent with an essential role of ACSL-dependent FA activation for entry of scavenged FAs into metabolic networks. These data therefore indicate that FA consumption occurs independent of perturbations to endogenous FA synthesis and, in contrast, can be directly blocked by disruption of proteins involved in the scavenging pathway.

As we previously demonstrated that lipid scavenging is essential for proliferation, we next tested the consequence of GSK and/or TriC treatment on proliferation kinetics. Consistent with the dominant role of lipid scavenging in supporting cell proliferation, treatment with GSK had no effect on the proliferation kinetics of H1299 NucRFP cells, while TriC treatment impaired cell proliferation (Fig 4E). Notably, when lipid uptake capacity was constrained by TriC treatment, GSK co-treatment now caused a modest additional antiproliferative effect. These data indicate that, while fatty acid synthesis capacity appears unable to meet lipid synthesis demands on its own, it can modestly contribute to proliferation when lipid uptake is constrained. Finally, to assess whether the relationship between lipid synthesis and scavenging is conserved across various cell lines, we tested the effects of GSK and/or Tri C on 143B, CCLP1, A549, HCT116, and HT1080 cells grown in the serum-free media by cell counts (Fig 4F). As with H1299 cells, all cell lines showed sensitivity to TriC treatment and were insensitive to GSK treatment alone. Interestingly, however, there were heterogeneous responses to the combination of TriC and GSK, with some cell lines showing additional proliferation defects upon GSK co-treatment (H1299, 143B, CCLP1) while others were unaffected (A549, HCT116, HT1080), presumably reflecting cell intrinsic differences in lipid synthesis capacity (Fig 4E-F). Collectively, these findings demonstrate that lipid scavenging is essential for proliferation and that *de novo* lipid synthesis is dispensable in the presence of exogenous lipid sources.

## Discussion

FBS, a ubiquitous additive in culture media, supports proliferation across numerous cell types; however, the specific serum components needed for growth are not well defined. Here, we combined live-cell imaging with LC-MS to show that metals and albumin-associated lipids in serum are crucial factors for sustaining cancer cell proliferation in culture. We observed that albumin-bound lipids, such as FA, LPC, and LPE, were rapidly consumed from the media, while lipoprotein-transported lipids were minimally depleted during proliferation. Furthermore, we found that a defined mixture of insulin, transferrin, selenium, ethanolamine and various trace metals was an effective serum replacement (predominantly for providing metals) when combined with albumin-associated lipids, providing a tool to investigate lipid consumption in a controlled lipid environment. Using this defined media, our results support a model whereby albumin-associated lipid consumption occurs in a mass-action manner independent of FA structure, with minimal competition with other lipid classes. This uptake did not correspond to lipid demand, as inhibiting *de novo* lipid synthesis had minimal effects on lipid scavenging, however direct disruption of components in the scavenging pathway blocked lipid uptake and impaired cell proliferation.

Our results contribute to the development of serum-free tissue culture media, which could have benefits for research beyond the cost savings and ethical considerations of omitting FBS (Piletz et al. 2018). For instance, recent data has shown that many genetic dependencies are influenced by the nutrient composition in the microenvironment; therefore serum-free conditions provide an opportunity to uncouple the numerous metabolic variables inextricably supplied by FBS (Rossiter et al. 2021; Lagziel, Lee, and Shlomi 2019). Although it could be argued that the molecular complexity of FBS more closely models *in vivo* conditions, the composition and abundance of bioactive molecules may also be inappropriate for modelling the cellular microenvironment experienced in an adult human and thus may mask important phenomena. In addition, our work is in agreement with other related studies in finding that FBS lots exhibit substantial variation in their metabolomic profiles, particularly lipids, which also significantly differs from human serum (Liu et al. 2023; Else 2020). This work is thus a step towards defined media conditions, which may facilitate future studies using genetic screens in cells grown in defined media to uncover which metabolic and signaling pathways are buffered by serum constituents, thereby potentially identifying additional serum factors that support proliferation in other contexts.

One of the major findings from using a serum-free media formulation to explore lipid consumption is that FA scavenging in proliferating cells is primarily driven by the concentration of exogenous lipids rather than by selective uptake based upon lipid structure. Indeed, treatment of cells with increased levels of FAs is associated with several effects on cell physiology including driving lipid droplet formation, causing lipotoxicity when excess saturated FAs accumulate intracellularly, and promoting ferroptosis when excess polyunsaturated FAs are enriched in membrane lipids (Akiyama et al. 2023; Antunes et al. 2022; Lipke, Kubis-Kubiak, and Piwowar 2022; Safi, Menéndez, and Pol 2024). The vulnerability to these outcomes based on lipid environment thus supports our conclusion that FA consumption rate is primarily determined by exogenous availability rather than metabolic need, making cells susceptible to the abundance and composition of environmental lipids. This results therefore underscore the potential utility of nutritional approaches that may modify the tumor lipid microenvironment to promote cancer cell lipid states that may be favorable for therapeutic targeting.

While proteins involved in FA and LPC transport (e.g. CD36 and MFSD2a) have been implicated to exhibit substrate preference based upon FA structure, it is unclear if this leads to relevant differences in substrate consumption during cell proliferation (Terry et al. 2023; Nguyen et al. 2014). Consistent with the minimal selectivity of scavenging based on fatty acyl structure, screens aimed at identifying lipotoxicity modifiers have predominantly uncovered factors that regulate intracellular lipid homeostasis, such as those involved in lipid droplet formation and phospholipid remodeling, rather than lipid transporters (Piccolis et al. 2019; Magtanong et al. 2019). Moreover, because lipid consumption is proportional to exogenous concentrations, interpretations of selective lipid uptake may instead reflect differences in initial abundance and lipid stability.

Despite the increase in lipogenic enzyme expression commonly observed in cancer cells, our data underscore an indispensable role of exogenous lipids in fulfilling the lipid requirement for proliferation (Röhrig and Schulze 2016; Szutowicz, Kwiatkowski, and Angielski 1979). Indeed, we observed exogenous lipid sources sustained serum-free cultivation of cells, and our results are consistent with the minimal effects of disruption to *de novo* lipid synthesis on cell proliferation when exogenous lipids are available (Kuhajda et al. 1994). The critical nature of lipid scavenging is further supported by the modest clinical efficacy of FASN inhibitors as a monotherapies for cancer treatment, as lipid-rich microenvironments *in vivo* may enable efficient lipid scavenging that diminishes the importance of *de novo* lipid synthesis (Wang et al. 2022; Lien et al. 2021; Medes, Thomas, and Weinhouse 1953). Indeed, cells cultured in physiologically relevant concentrations of lipids, which is approximately an order of magnitude greater than what is present in media containing 10% FBS, have been shown to preferentially incorporate scavenged lipids into cellular membranes during proliferation (Yao et al. 2016; Else 2020). Further, we identify the ACSL family of proteins as crucial mediators of exogenous FA scavenging through metabolic trapping. Indeed, disruption of FA activation with triacsin C abolishes FA scavenging, consistent with the essentiality of FA condensation with coenzyme A for entry into the cellular lipidome (Mashek, Li, and Coleman 2007). Importantly, previous studies using genetic and pharmacological perturbations to ACSL family members indicate these proteins are crucial in FA incorporation into neutral lipids and phospholipids; our data additionally directly implicates this family of proteins in the FA scavenging pathway (Igal, Wang, and Coleman 1997). Interestingly, we find that only when FA scavenging was constrained could FA synthesis detectably contribute to cell proliferation, albeit heterogeneously across cell lines. Together, these findings suggest that targeting lipid scavenging could be a more effective therapeutic approach to target the increased lipid demand of proliferating cancer cells, though any such strategy would have to overcome the redundancy of alternate albumin-bound lipid sources.

## Acknowledgements

This research was supported in part through the NIH/NCI Cancer Center Support Grant P30 CA015704 through a pilot award and support for the Proteomics & Metabolomics Shared Resource of the Fred Hutch/University of Washington Cancer Consortium. L.B.S. acknowledges support from the NIGMS R35 GM147118.

