## Supplementary Figures for "Defined media reveals the essential role of lipid scavenging to support cancer cell proliferation"

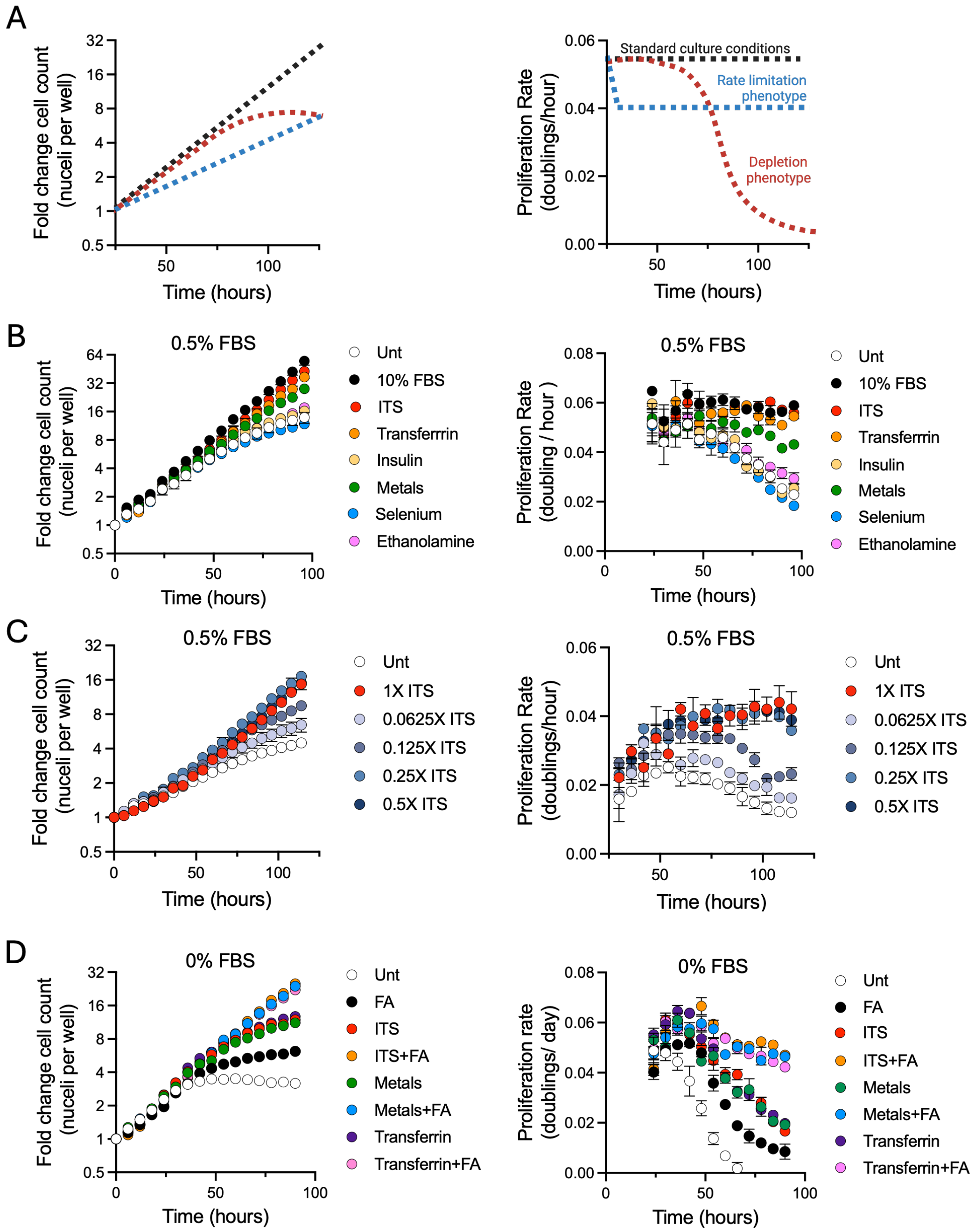


**Supplementary Figure 1. Serum-restriction results in a depletion phenotype.**
**(A)** Predicted changes to cell growth kinetics upon two theoretical modes of proliferation inhibition. Fold change in cell counts (left) and moving average of proliferation rate (right). The blue dotted line reflects a rate limitation phenotype (consistently slowed) compared to the red dotted line which reflects a depletion phenotype (progressive loss of proliferation over time), which can appear identical if only counting initial and final timepoints. **(B)** Growth kinetics of cells cultured in 0.5% FBS with individual components of the ITS mix. Fold change in cell counts (left) and moving average of proliferation rate (right). **(C)** Growth kinetics of cells cultured in 0.5% FBS supplemented with a decreasing titration of the complete ITS mix. Fold change in cell counts (left) and moving average of proliferation rate (right). **(D)** Growth kinetics of cells cultured in DMEM supplemented with and without FA, and either ITS, transferrin, or trace metals. Fold change in cell counts (left) and moving average of proliferation rate (right). Error bars represent mean ± SEM (n = 3).

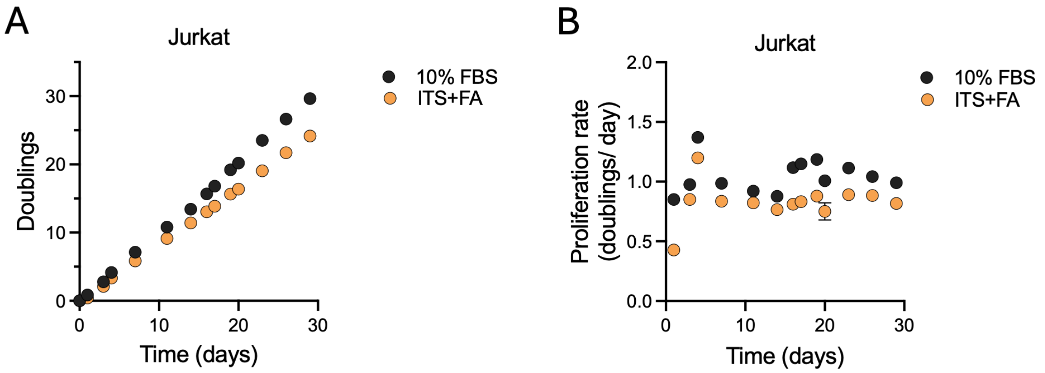


**Supplementary Figure 2: Serum free media supports sustained Jurkat cell growth.**
**(A-B)** Jurkat cell proliferation in serum-free media relative to culture media containing 10% FBS. Population doublings (A) and proliferation rate (B). Error bars represent mean ± SEM (n=3).

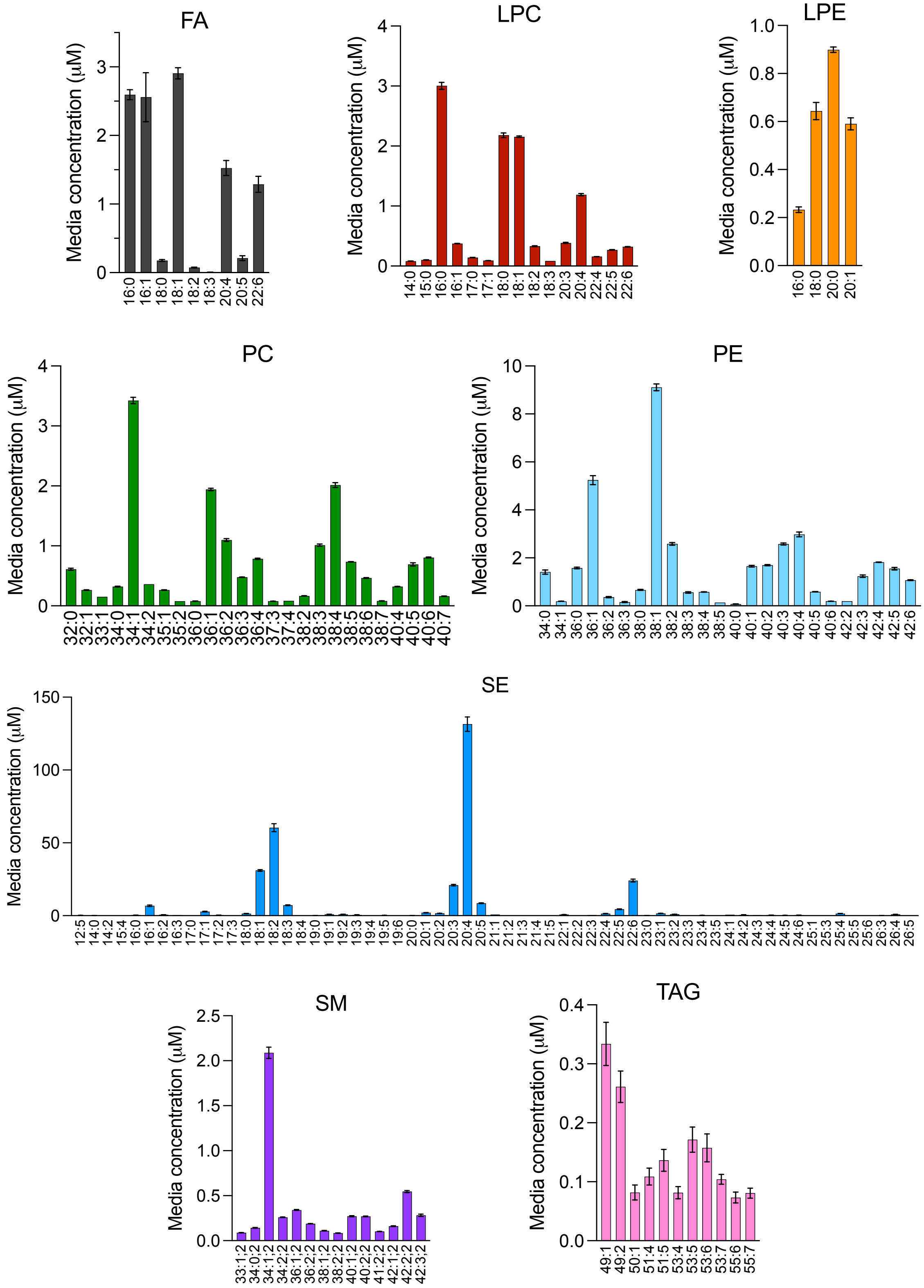


**Supplementary Figure 3. Lipid profiles of FBS.**
Lipid species that constitute the top 99% of the total lipid pool detected in 10% FBS containing media. The x-axis indicates the number of carbons and the number of double bonds present in each lipid species. Error bars represent mean ± SEM (n = 3). Abbreviations: SE, sterol esters; PE, phosphatidylethanolamine; PC, phosphatidylcholine; SM, sphingolipids; TAG, triacylglycerols; LPC, lysophosphatidylcholine; LPE, lysophosphatidylethanolamine; FA, fatty acids.

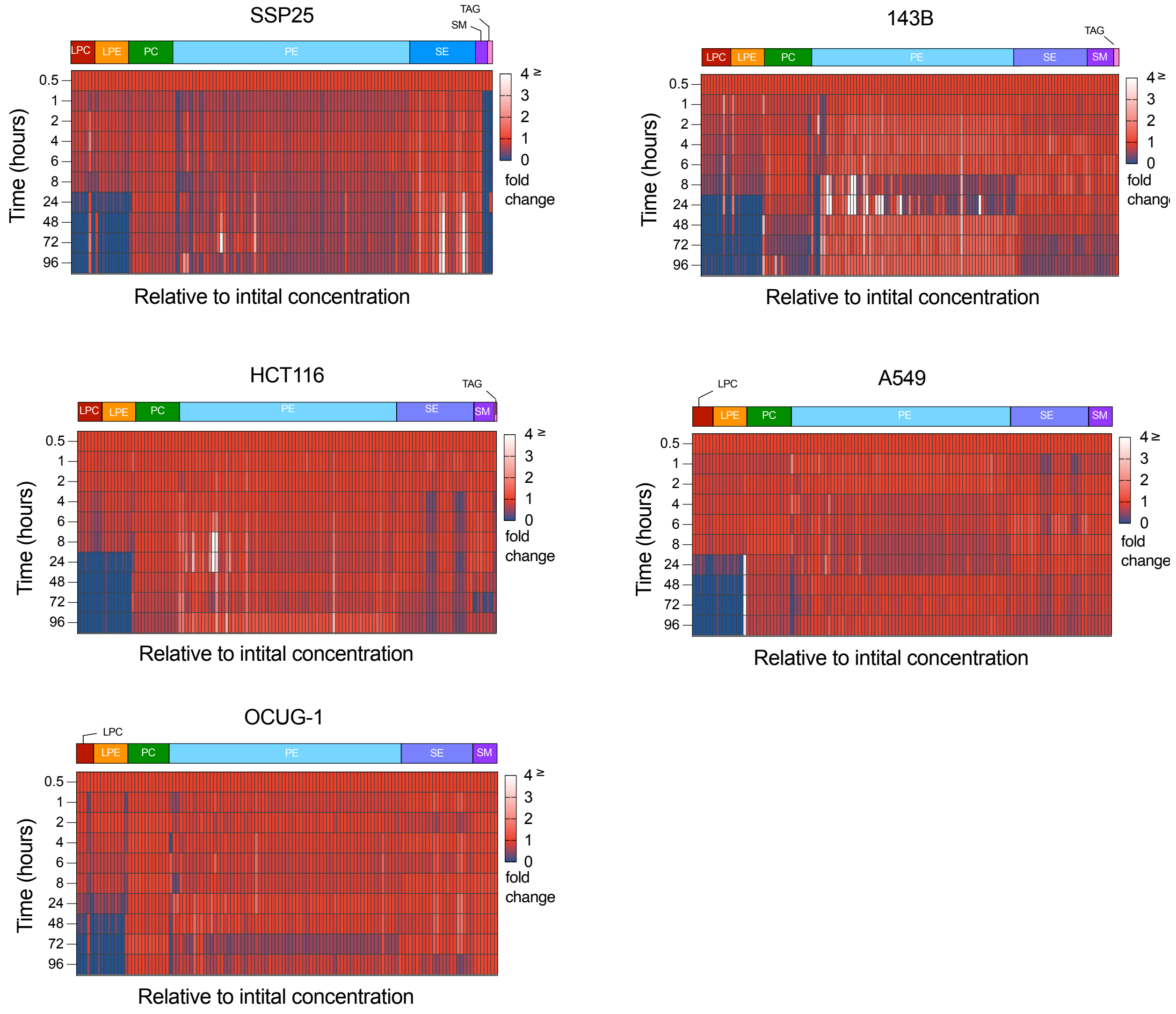


**Supplementary Figure 4. Lipid consumption patterns are conserved across cell lines.** Media lipid depletion from 10% FBS containing media for cholangiocarcinoma (SSP25), osteosarcoma (143B), non–small cell lung cancer (A549), colon cancer (HCT116), and gallbladder carcinoma (OCUG-1).
